# The large GTPase Guanylate-Binding Protein-1 (GBP-1) promotes mitochondrial fission in glioblastoma

**DOI:** 10.1101/2024.05.08.593235

**Authors:** Ryan C. Kalb, Geoffrey O. Nyabuto, Michael P. Morran, Swagata Maity, Jacob S. Justinger, Andrea L. Nestor-Kalinoski, Deborah J. Vestal

**Affiliations:** Department of Biological Sciences, University of Toledo, 2801 W. Bancroft St., Toledo, OH, 43606, USA; Department of Surgery, University of Toledo, 3000 Arlington Ave., Toledo, OH,43614, USA

**Keywords:** Dynamin-like Proteins (DLPs), Epidermal Growth Factor Receptor (EGFR), glioblastoma multiforme (GBM), Guanylate-Binding Protein-1 (GBP-1), immunofluorescence, mitochondrial dynamin-related protein 1 (Drp1), Translocase of Outer Mitochondrial Membrane 40 (TOMM40)

## Abstract

Glioblastomas (aka Glioblastoma multiforme (GBMs)) are the most deadly of the adult brain tumors. Even with aggressive treatment, the prognosis is extremely poor. The large GTPase Guanylate-Binding Protein-1 (GBP-1) contributes to the poor prognosis of GBM by promoting migration and invasion. GBP-1 is substantially localized to the cytosolic side of the outer membrane of mitochondria in GBM cells. Because mitochondrial dynamics, particularly mitochondrial fission, can drive cell migration and invasion, the potential interactions between GBP-1 and mitochondrial dynamin-related protein 1 (Drp1) were explored. Drp1 is the major driver of mitochondrial fission. While GBP-1 and Drp1 both had punctate distributions within the cytoplasm and localized to regions of the cytoplasmic side of the plasma membrane of GBM cells, the proteins were only molecularly co-localized at the mitochondria. Subcellular fractionation showed that the presence of elevated GBP-1 promoted the movement of Drp1 from the cytosol to the mitochondria. Migration of U251 cells treated with the Drp1 inhibitor, Mdivi-1, was less inhibited in the cells with elevated GBP-1. Elevated GBP-1 in GBM cells resulted in shorter and wider mitochondria, most likely from mitochondrial fission. Mitochondrial fission can drive a number of important cellular processes, including cell migration, invasion, and metastasis.

**Simple Summary:** Glioblastomas are the most common and most aggressive adult brain tumors arising from astrocytes. Up to 60% of glioblastomas are promoted by either amplification and/or mutation of the Epidermal Growth Factor Receptor (EGFR). The large GTPase Guanylate-Binding Protein-1 (GBP-1) is one of the most robustly induced proteins following EGFR signaling in glioblastomas. How GBP-1 promotes tumor progression in glioblastomas is still under investigation. This study shows that GBP-1 promotes tumor progression by associating with Drp1 at glioblastoma mitochondria and enhancing their fission. Specifically, increased mitochondrial fission can promote tumor cell migration, invasion, and metastasis.

## Introduction

Glioblastomas (aka Glioblastoma multiforme (GBM)) are the highest grade of adult brain tumors arising from astrocytes, a class of neuronal support cells in the brain [1]. GBMs are the most abundant and most aggressive adult brain tumors [2,3]. This aggressiveness results, in large part, from their high levels of both proliferation and invasiveness [4]. Even after tumor resection, radiation, and chemotherapy, the median survival rate is about 14-15 months [5].

Epidermal Growth Factor Receptor (EGFR) signaling plays an important role in the prognosis of GBMs. Up to 60% of GBMs have either amplified wild type EGFR or mutant EGFRvIII (amplified or not) driving their proliferation and/or invasion [6,7]. This would suggest that anti-EGFR therapy would be a good approach for treatment of GBMs but unfortunately anti-EGFR treatments have not proven efficacious in GBMs. As a consequence, some attention has shifted to identifying critical downstream proteins of EGFR signaling to target. One such protein is the large cytokine-induced GTPase, Guanylate-Binding Protein-1 (GBP-1) [8–11].

GBP-1 is part of a large cytokine-induced protein family that are members of the dynamin superfamily of GTPases [12–16]. GBPs differ from most GTPases by hydrolyzing GTP to both GDP and GMP, which influences whether the proteins are monomers, dimers, or polymers [17–29]. GBP-1 is the best studied of the GBP family [21,29,30]. While most commonly studied for their role in immune responses, the study of GBPs has also focused on their complicated role in cancer development and progression [31–33]. The anti-tumor activity of GBP-1 was first highlighted by the finding that cytokine-induced inhibition of angiogenic growth factor-[34]induced proliferation of cultured human endothelial cells resulted in the expression of GBP-1 [35,36]. Forced expression of GBP-1 in HUVECs inhibited their proliferation [35]. GBP-1 also inhibited the invasion and angiogenic ability of endothelial cells by inhibiting the expression of MMP-1 *in vitro* [36]. It was also associated with decreased angiogenesis *in vivo* [36]. Additional data on whether GBPs are protective or promote tumorigenesis is conflicting [14,37,38]. The data suggests that the roles of GBPs in cancers depend on such variables as cell type and immunological environment [39]. Data suggests that some of the effects of GBP expression may be the consequence of whether the tumor is in a primarily interferon driven environment versus a growth factor driven one.

Several labs have explored the role of GBP-1 in GBMs [8–11]. In the first study, U87 and U178 GBM cells were transduced with EGFR, followed by incubation with or without EGF for 3 hours and gene array analysis to identify the most abundant EGF-induced proteins in GBMs [8]. The most robustly induced genes after EGF treatment were associated with interferon (IFN) signaling, including GBP-1 [8]. Twelve out of 19 human GBM tumor samples had elevated GBP-1 mRNA, while EGFR mRNA was elevated in 9 of them [8]. To extend analyses, 8 GBM tumors with corresponding normal brain samples were examined. EGFR levels were higher in all 8 tumor samples compared to paired normal brain. GBP-1 levels were higher in 7 out of 8 patients. Matrix metalloproteinase-1 (MMP-1) expression was elevated in 4 of the eight tumors compared to normal tissue. Two of the tumors had elevated GBP-1 expression with no MMP-1. Ten of the 10 GBM cell lines examined all expressed significant EGFR and GBP-1 without EGF treatment, but only 4 expressed MMP-1^1^ [8]. The observation that 4 of the 8 tumors and 4 of the 10 cell lines expressed MMP-1 prompted exploration of a possible connection between GBP-1, EGFR, and MMP-1. Knockdown (KD) of GBP-1 in U87 cells over-expressing EGFR resulted in attenuated ability of EGF treatment to induce the expression of MMP-1 [8]. It also reduced the EGF driven invasion *in vitro* [8]. Forced expression of GBP-1 in A1207 cells promoted cell invasion that was attenuated by si-RNA KD of MMP-1. This prompted the conclusion that GBP-1 is required for the EGF induction of MMP-1 [8]. Interestingly, KD of GBP-1 in SNB19 GBM cells resulted in significantly less tumor invasion than cells with control shRNA after intracranial xenograft [8]. Forced expression of GBP-1 in both SHG44 and U251 GBM cells caused a modest but statistically significant increase in cell migration and more significant increase in cell invasion and overexpression of GBP-1 in SHG44 cells promoted tumor growth and shortened survival [11]. In fact, GBP-1 is an independent risk factor of prognostic value for GBMs [11].

GBP-1 was also implicated in GBM development/progression subsequent to analysis of signaling networks downstream of EGFR and EGFRvIII in GBM xenografts [10]. Xenografts were generated from 8 human GBM tumors and analyzed for EGFR expression by western blot (WB). Two of the xenografts expressed wild type EGFR without amplification, three expressed amplified EGFR, and 3 expressed amplified EGFRvIII [10]. Comparison of xenografts containing amplified EGFRvIII compared to xenografts with either unamplified or amplified levels of wildtype EGFR identified 4 genes specifically upregulated in the EGFRvIII cells. These were GBP-1, SA100A10, CAIII, and MVP [10]. Only CAIII was amplified in all 3 of the EGFRvIII cells. Kaplan Meier analysis showed that elevated levels of each of the three genes resulted in poorer percent survival [10]. Why this xenograph study did not show elevation of GBP-1 mRNA in the xenographs with wild type EGFR remains unclear. To further understand how GBP-1 affects GBM cells, forced expression of EGFRvIII in U87 cells (U87-EGFRvIII) promoted cell proliferation and KD of GBP-1 in U87-EGFRvIII cells significantly inhibited tumor growth *in vitro* [9]. EGFRvIII overexpression in U87 cells promoted tumor growth in subcutaneously inoculated mice compared to parental cells, and KD of GBP-1 significantly inhibited tumor growth [9]. Forced expression of GBP-1 in U87-EBFRvIII cells promoted tumor development but did not promote cell proliferation *in vitro* [9]. Elevated expression of GBP-1 promoted cell invasion in xenograft intracranial injections and resulted in much larger tumor volumes. This elevated growth and invasion resulted in significant reduction in percent survival [9]. Forced expression of EGFRvIII in U87 GBM cells showed significant upregulation of GBP-1 [9]. Upregulated GBP-1 mRNA was correlated/associated with EGFRvIII expression in 20 GBM tumor samples [9].

In an effort toward determining the mechanism(s) by which GBP-1 promotes cell migration/invasion, the intracellular localization of GBP-1 in GBM cells was analyzed. Surprisingly GBP-1 strongly localized to mitochondria in GBM cells by both confocal microscopy and subcellular fractionation. Specifically, GBP-1 localized to the cytosolic side of the outer mitochondrial membrane of mitochondria. GBP-1 expression resulted in the relocation of the mitochondrial dynamin-related protein 1 (Drp1) to the mitochondria where GBP-1 and Drp1 co-localize. Forced expression of GBP-1 results in shorter, wider mitochondria. This was intriguing since mitochondrial dynamics are associated with regulation of migration and invasion [40]. Our study shows that GBP-1 localizes to mitochondria in GBM cell lines and can play a role in regulating their mitochondrial behavior to promote the shortening of the mitochondria.

## Materials and Methods

### Cells and cell culture

U251 and SNB75 glioblastoma cells were obtained from American Type Culture Collection (ATCC) and maintained in complete DMEM (Mediatech, Manassas, VA, USA) with 4.5g/L glucose was supplemented with 10% fetal bovine serum (FBS; Atlanta Biologicals, Lawrenceville, GA, USA), 2mM L-glutamine (Mediatech), and 50 µg/ml penicillin/streptomycin (Mediatech)). Stable cell lines containing control vector pIRES-hygro2 and high expressing hGBP-1 c-myc-hGBP1-pIRES-hygro2 were maintained in complete DMEM with the addition of hygromycin (50 µg/ml) (Research Products International, Mount Prospect, IL, USA). All cells were cultured at 37°C and five percent carbon dioxide. Cells were suspended by treatment with 0.05% trypsin/0.53 mM EDTA (Mediatech). Cells were regularly checked for mycoplasma, which will induce interferons and alter the GBP-1 expression in “untreated” cells.

### PAGE gels and Western Blot

Cells at 80-85 percent confluence were lysed in RIPA buffer (50 mM Tris pH 7.5, 150 mM NaCl, 1% NP40, 0.5% sodium deoxycholate, and 0.1% SDS) containing 1mM phenylmethylsulfonyl fluoride and one percent protease cocktail (Sigma). Cell lysates (20 µg) were size fractionized by 8% polyacrylamide SDS-PAGE gels, transferred onto Immobilon-P membranes for 2 hours at 100 volts in 1x transfer buffer (25 mM Tris, 0.192 M Glycine, 20% methanol) for western blot analysis as previously described [34]. The membrane was blocked in blocking buffer (TBST (250 mM Tris, pH 8, 5 M NaCl, 0.3% Tween-20) plus 5% nonfat dry milk) for 1 hour at room temperature or overnight at 4°C. After transfer, membranes received primary antibody in blocking buffer for 1.5 hours. The primary antibodies were rat anti-hGBP-1 (1b1) (Santa Cruz Biotechnology 1:10,000); mouse anti-MMP1 Mab 901 (R&D Systems 1:750); rabbit anti-actin (Sigma 1:1000), rabbit anti-Translocase of Outer Mitochondrial Membrane 40 (TOMM40; (1:1500)), mouse anti-myc (1:2000), and mouse anti-Drp1 (1:100, Santa Cruz sc-101270). Membranes received secondary antibody diluted in blocking buffer for at least 1 hour. The hrp-conjugated secondary antibodies were: goat anti-rabbit IgG (1:1000; Invitrogen), goat anti-rat IgG (1:1500; Rockland), goat anti-mouse IgG (1:1000; Invitrogen). Antibodies were detected using Super Signal West Pico Chemiluminescent Substrate (Thermo) following manufacturer’s instructions.

### Immunofluorescence & Microscopy

Cells (50,000) were plated per cover slip in a 6 well dish and incubated for 24 hours in complete DMEM (Mediatech). Cells were fixed with 4% paraformaldehyde (PFA) for 20 minutes and washed in phosphate buffer saline (PBS; 138 mM NaCl, 2.6 mM KCl, 5.4 mM Na2HPO4, 1.8 mM KH2PO4, pH 7.4). An antigen retrieval step was used only with primary anti-GBP-1 antibody 1b1 or any combination of antisera that contained 1b1. Cover slips were placed in antigen retrieval buffer (10 mM TRIS, 1.3 mM EDTA, 1L H_2_O, pH 9), and submerged in a hot water bath at 95°C for 10 minutes. Cells were then permeabilized with 0.1% Triton X-100 in PBS for 10 minutes and blocked with antibody dilution buffer (10% PBS, .05% Tween-20, 6% BSA, 5% Glycine) with 10% non-immune horse serum for 2 hours. Cells were incubated with primary antibodies overnight at 4°, followed by PBS washes and addition of secondary antibodies for 1 hour at room temperature. Antibodies against TOMM40 (1:100; Proteintech (18409-1-AP)) and Cytochrome c (1:50; Abcam) were used for mitochondrial staining. Anti-GBP-1 (1b1) (1:25; Calbiochem or Santa Cruz), anti-myc (1:50; Proteintech (cat # 60003-2)) were used for hGBP-1. Anti-Drp1 (1:750; sc-101270) were from Santa Cruz. DAPI (4′,6-diamidino-2-phenylindole; 75 nM) was used for nuclear staining. The primary antibodies were detected by incubation with highly cross-absorbed Alexa 488-conjugated anti-rabbit (1:500), Alexa 488-conjugated anti-mouse (1:5,000), Alexa 594-conjugated anti-rat (1:500) and Alexa 594-conjugated anti-mouse (1:500). Cells were washed with PBS and stained with 150 nM DAPI for 5 min at RT. Coverslips were mounted with Fluoromount-G (SouthernBotech). Epifluorescent images were captured using an EVOS FL inverted microscope by American Microscope Group (AMG) (Thermo Fisher Scientific) with a 40X oil objective and GFP, Texas Rd, Cy5, and DAPI filters. Confocal images were collected using a TCS-SP 5 spectrophotometric multiphoton laser scanning confocal microscope (Leica Microsystems) collecting 0.2 - 1 µm optical sections.

### Mitochondrial Isolation

Mitochondria were isolated from GBM cells using the Mitochondria Isolation Kit for Cultured Cells (Thermo Scientific) following manufacturer’s directions. Cells (1.5 – 2 × 10^7^) were grown in 150 mm dishes, scraped and their pellets were frozen at −80°C. The pellet from a single 150 mm dish was used for each mitochondrial isolation. Briefly, after cell lysis the nuclei are removed by a low-speed centrifugation. The total cell lysate (TCL) aliquot is taken at this time. Further centrifugations will separate the mitochondrial fraction from the cytosol. Alter lysis of the mitochondria, the protein levels of the TCL, cytosol, and mitochondrial fraction were determined and equal amounts of protein from each fraction were separated by SDS-PAGE and analyzed by western blot as described above.

### Generation of amino-terminal myc-tagged GBP-1 in pIRES-hygro2

The generation of pCMV2(NH) Flag-hGBP-1 was described previously [41]. Next an amino terminal myc-epitope tag was added to pIRES2-eGFP. pIRES2-eGFP was digested with *BglII* and *EcoR1* and purified. The myc epitope was generated by the hybridization of two oligonucleotides: forward 5’-gatctatggaacagaaactgatcagcgaggaagatctgaatcgcggccgcc-3’ and reverse 5’-aattcgcggccgcgattcagatcttcctcgctgatcagtttctgttccata-3’. When the oligos are annealed, the 5-end has an overhang compatible with a *BglII* site and the 3’ end is compatible with an *EcoRI* site. Ligation of the double stranded oligonucleotide generated c-myc pIRES2-eGFP. To generate c-myc hGBP-1 pIRES2-eGFP, c-my pIRES2-eGFP was digested to completion with SmaI and then partially digested with *NotI*. The appropriate fragment was gel purified and ligated with purified hGBP-1 cDNA liberated from pCMV/flag/NH GBP-1 by SmaI/NotI digested. To generate c-myc hGBP-1 pIRES-hygro2, pIRES-hygro2 was cut to completion with *NheI/SmaI* and ligated to the insert released from c-myc hGBP-1-pIRES2-eGFP by *NheI/SmaI*. The construct was sequenced to assure fidelity.

### Generation of U251 cells expressing myc-tagged hGBP-1

U251 cells were transfected with c-myc-hGBP1-pIRES-hygro2 (2 µg) or pIRES-hygro2 (2 µg). Transfection solutions were made at a lipid (µg) to plasmid (µL) ratio of 3:1 using FuGene 6 (Promega) as the transfection reagent. Transfected U251 cells were selected in complete media with 50 µg/ml hygromycin. After 6 weeks of expansion from 12 well dishes to 10 cm dishes, twelve of the c-myc-hGBP1-pIRES-hygro2 transfected U251 colonies were analyzed by Western Blot for expression of hGBP-1. For further studies the cells chosen were (control) A10, and (GBP-1 expressing cells) #4 and #7.

### Mitochondrial measurement

To measure mitochondrial length and width, cells were stained for TOMM40 and imaged with a Leica TCS SP5 multiphoton confocal microscope at 0.2 µm optical sections. Following 3D reconstruction, the .lif files were opened in LASX and 10 mitochondria were measured from each of 20 randomly chosen cells for each cell type/condition. The data from two experiments were combined and analyzed using box and whisker plots from GraphPad Prism at 99% confidence interval. P values were determined by students t-test.

### Wound healing assay

SNB75 cells, U251 cells, and U251 cells containing myc-tagged GBP-1 were grown to confluence in flat bottom clear 96-well plates. Visual confluence was monitored for all seeded wells utilizing the IncuCyte S3 Live Cell Analysis System (Sartorius). Prior to scratch, complete growth media was removed and replaced with serum free complete media for 4 hours. After serum starvation, cells were “scratched” with a 96-well WoundMaker TM (Sartorius) and washed twice with sterile PBS to remove any suspended cells or debris caused by the scratch. Complete growth media was applied to all wells containing either DMSO or Mdivi-1 (Selleckchem Cat#S7162) resuspended in DMSO at 5 µM, 20 µM, or 50 µM. Mdivi-1 is the most frequently used Drp1 inhibitor but it has some limitations [42]. Mdivi-1 can inhibit oxidative metabolism and modestly inhibit antioxidant activity [43,44]. Scratch closure was tracked every 2 h for 24 h via the IncuCyte S3 Live Cell Analysis System at 10X magnification. Wound width data analysis was generated for all wells utilizing the IncuCyte Scratch Wound analysis software module. Raw wound width data was exported and converted into wound % closure values for each individual sample with n=5 per treatment group.

### Statistics

Statistical analyses were carried out by two-tailed t-tests when two groups were analyzed. Statistically different groups are defined as * = p <0.05, ** = p <0.01, *** = p < 0.001.

## Results

### EGF treatment does not induce GBP-1 in all cultured GBM cells

To determine the universality of EGFR signaling inducing GBP-1, GBM cell lines were treated with EGF or IFN-γ and analyzed for the induction of GBP-1 (Fig. 1A). IFN-γ was used to confirm that the cells were able to make GBP-1. Each membrane contained U251 cells so that the relative intensity of GBP-1 could be visualized. Interestingly, only four of the cell lines expressed little or no GBP-1 before treatment (U251, SNB19, U87, LN229). EGF treatment modestly induced GBP-1 in only 2 of these (U87 and SNB19). GBP-1 was constitutively expressed in SNB75, SNB295, and T98G cells. It was modestly induced in SNB295 cells. GBP-1 levels were reduced in T98G cells with EGF treatment but remained the same in SNB75 cells (Fig. 1A). All cells responded to IFN-γ by inducing GBP-1 (data not shown)^(Footnote1)^. Previous studies indicated that the induction of GBP-1 by EGFR signaling promoted invasion by inducing the expression of MMP-1 [8]. Surprisingly, only SNB75 cells responded to EGF treatment by inducing MMP-1 (Fig. 1B). MMP-1 was modestly expressed in SNB295 cells prior to treatment but expression was lost after EGF or IFN-γ treatment (not shown). MMP-1 was not expressed in any of the cells after treatment with IFN-γ. This data (along with information presented in the introduction) were interpreted to suggest that EGF-mediated induction of MMP-1 expression via GBP-1 and its promotion of migration are most likely not the only mechanism to promote migration and invasion of GBM cells by GBP-1.

**Figure 1.**
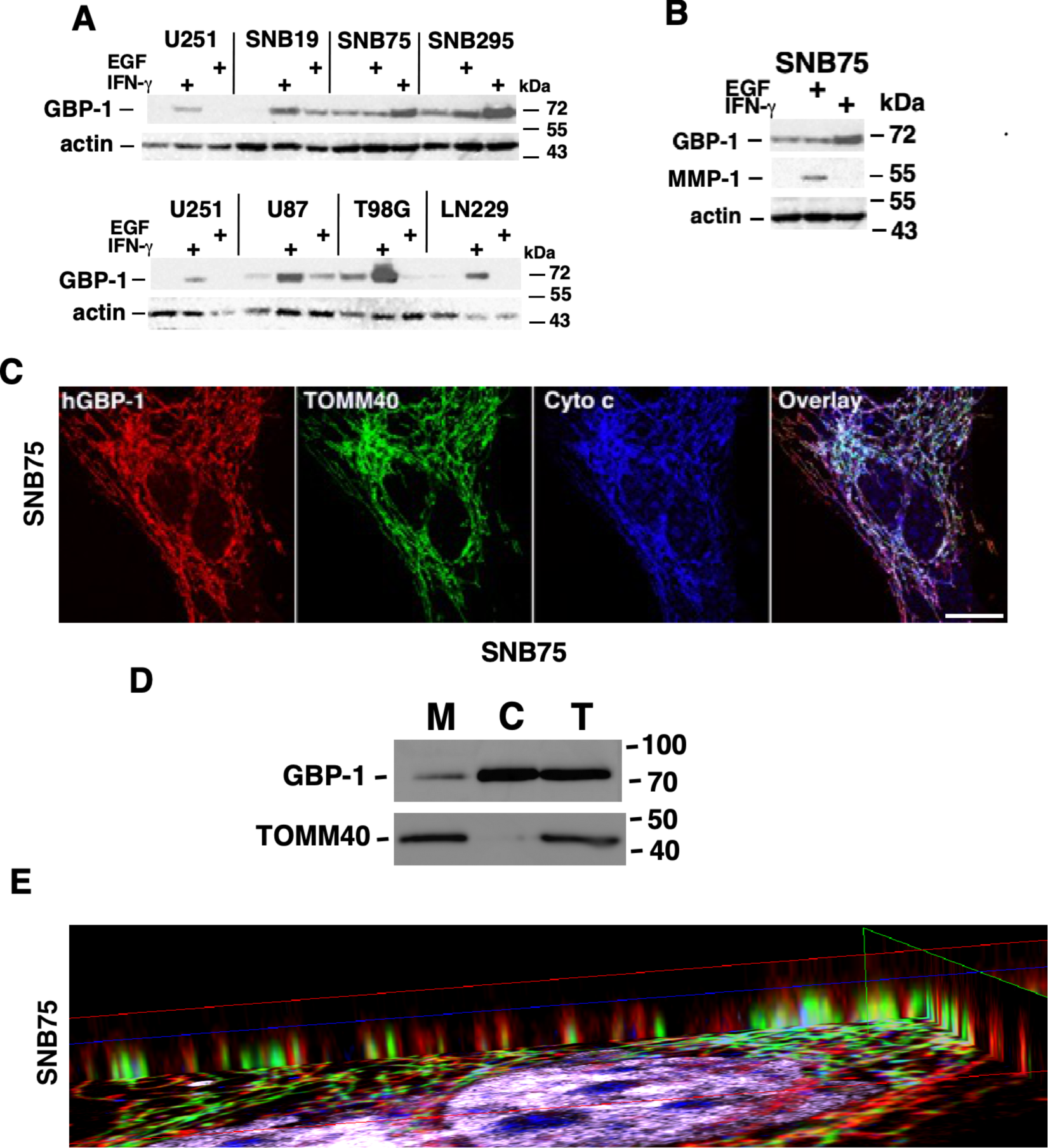
GBP-1 localizes to mitochondria in SNB 75 glioblastoma cells by indirect immunofluorescence. **A.** GBM cell lines were plated overnight and then serum starved for 24 hours. Cells were either left untreated or treated with 50 ng/ml hEGF or 500 U/ml of hIFN-γ for 24 hours. Cell lysates (20 µg) were separated by 8% SDS-PAGE, followed by western blot for GBP-1 and actin. U251 was run into each membrane as a visual marker for representative intensities. **B.** WB of SNB75 was shown with lane for MMP-1 expression. **C.** SNB75 cells were analyzed by triple label IF to determine the intracellular location of GBP-1. After incubating with antibodies against GBP-1, TOMM40, and cytochrome c, cells were incubated with appropriate highly cross adsorbed secondaries, mounted, and images were capture by confocal microscopy at 1 µm optical sections. Size bars = 25 µm. **D.** SNB75 cells were fractionated into mitochondrial (M) and cytosolic (C) fractions and analyzed for GBP-1 and TOMM40 by western blot. Total cell lysates (T) were also provided. **E.** SNB75 cells were analyzed by multiphoton confocal microscopy. The image series from one z-stack is shown in Fig 1C. SNB75 images were compiled and analyzed by 3D reconstruction. Visualization of the image from the side suggests that the TOMM40 signal (green) for the outer mitochondrial membrane is surrounded by GBP-1. This suggests that GBP-1 is on the outside of the outer mitochondrial membrane.

### GBP-1 localizes to the mitochondria of GBM cells

As a step toward narrowing the analysis of GBP-1 function, the intracellular localization of GBP-1 in SNB75 cells was analyzed by indirect immunofluorescence (IF) (Fig. 1C). Surprisingly, GBP-1 localized to elongated, filamentous structures that suggested mitochondria (Fig. 1C). Co-staining SNB75 cells for TOMM40 and cytochrome c identified these structures as mitochondria (Fig. 1C). Note that while all three proteins localize to the mitochondria, their distribution at the mitochondria is not completely overlapping (Fig. 1C overlay). This is consistent with the different known localizations of the proteins in mitochondria. TOMM40 localizes to the outer membrane of the mitochondria, as a component of the outer mitochondrial membrane transporter or Translocon [45]. Cytochrome c localizes to the mitochondrial intermembrane space, where it serves as an electron transporter in the electron transport chain [46].

To confirm that GBP-1 associates with mitochondria, SNB75 cells were subcellularly fractionated to isolate mitochondria (Fig. 1D). As expected TOMM40 localized to the mitochondria (M). A very small amount of TOMM40 could be found in the cytosolic fraction (C), which could be accounted for by the fact that the gene for TOMM40 is nuclear (National Library of Medicine, Gene ID: 10452. Updated 26-Jan-2024). GBP-1 localized to both the mitochondria and the cytosol (Fig. 1D). The intracellular localization of GBP-1 by indirect immunofluorescence (IF) suggests that most of the GBP-1 is associated with mitochondria in SNB75 cells but the subcellular fractionation suggests that most of the GBP-1 is associated with the cytoplasm. GBP-1 is a lipid anchored protein that can form polymers and can come on and off cellular membranes using the farnesyl lipid on its C-terminus [47–49]. The observation that after subcellular fractionation most of the GBP-1 localized to the cytosol suggests a loose association of GBP-1 with the mitochondria that can be easily dissociated.

### GBP-1 localizes to the outer membrane of mitochondria

To get a clearer idea of where GBP-1 resides on mitochondria, images of SNB75 cells captured in Fig. 1C were compiled and analyzed by 3D reconstruction. Visualization of the image from the side shows that the TOMM40 signal (green) for the outer mitochondrial membrane is surrounded by GBP-1 (red) (Fig. 1E). This indicates that GBP-1 is on the cytosolic side of the outer mitochondrial membrane, which would be consistent with it being a lipid anchored protein relatively loosely associated with the cytosolic side of mitochondria.

To confirm that the localization of GBP-1 to GBM mitochondria was not unique to SNB75 cells, U251 cells expressing myc-tagged GBP-1 were generated (Fig. 2A). The cells will be designated U251+GBP-1 cells in the future. Triple label IF again demonstrated that GBP-1 localizes to the mitochondria (Fig. 2B). Interestingly, the morphologies of the mitochondria were very different between the two cell lines. The mitochondria of SNB75 cells were much more elongated and tubular than those of the U251 cells, despite both cell lines expressing GBP-1 that localized to the mitochondria. The far-right panel of this figure shows an enlargement of the area within the square in the cytochrome c panel. Consistent with the staining of SNB75, while all three proteins are found at the mitochondria, they are in slightly different locations. Again, to get a clearer idea of where GBP-1 resides on mitochondria, images of U251+GBP-1 cells were compiled and analyzed by 3D reconstruction. While the morphology of the mitochondria in these cells was less tubular than in the SNB 75 cells, visualization of the image from the side again shows that the TOMM40 signal (green) for the outer mitochondrial membrane is surrounded by GBP-1 (red) (Fig. 2C), again indicating that GBP-1 localizes to the cytosolic side of the outer membrane of mitochondria.

**Figure 2.**
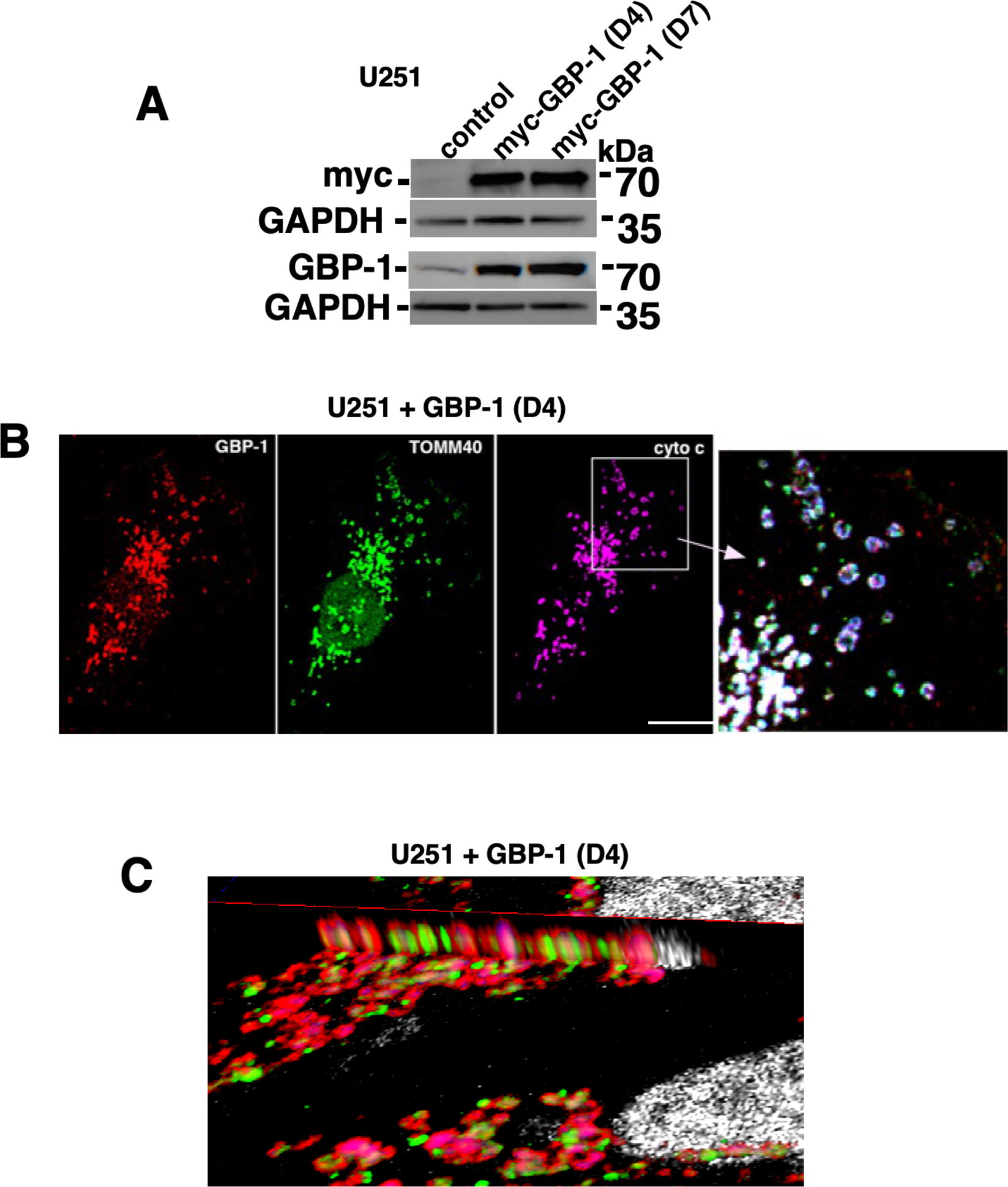
GBP-1 localizes to mitochondria in U251 cells. **A.** U251 cells were generated to express myc-tagged GBP-1. Control cells (empty vector) and two pools of cells expressing myc-tagged GBP-1 were analyzed by western blot for GBP-1 expression. **B.** U251+GBP-1 (D4) cells were analyzed by triple label IF for the localization of GBP-1 using antibodies against myc, TOMM40, and cytochrome c. The panel on the far right shows the overlay of the region of the photomicrograph designated by the box in the cytochrome c panel. Size bar = 20 µm. **C.** U251 cells expressing GBP-1 were analyzed by multiphoton confocal microscopy for the expression of TOMM40 and GBP-1. The images were compiled and analyzed by 3D reconstruction. Visualization of the image from the side suggests that the TOMM40 signal (green) for the outer mitochondrial membrane is surrounded by GBP-1 (red). This suggests that GBP-1 is on the outside of the outer mitochondrial membrane.

### GBP-1 did not promote migration in U251 cells, as measured by scratch assays

To determine whether our U251 cells responded to GBP-1 by enhancing migration, U251 and U251+GBP-1 cells were analyzed for migration by scratch assay (Supp. Fig. 1). U251 cells closed their wound 4 to 6 hours faster than U251 cells with GBP-1 (Supp. Fig. 1A). The wounds for U251 cells closed at about 14 hours after wounding and the U251+GBP-1 cells closed their wounds in about 20 hours. At least one wound closure of a second U251+GBP-1 (D7) cell line also closed their wounds at 20 hours (Supp. Fig. 1A). Closer examination of the wound closing curves shows that the presence of GBP-1 results in a flatter curve for the first 4 hours after wounding than observed in the absence of GBP-1 (delineated by the box) (Supp. Fig. 1A). This suggests about a 4-hour lag time before starting increased speed of migration after wounding. This 4 hour “lag” was also observed for the U251+GBP-1 (D7) cells (Supp. Fig. 1A). Representative images of wound closures are shown (Suppl. Fig. 1B). To determine the rate of wound closure, the slope of the graph of percent wound closure was determined from the closure data from 4 hours to 14 hours, chosen because of the “delay” in closure within the first 4 hours and because the wound was closed for U251 cells by 14 hours (Supp. Fig. 1C). The rate of wound closure was greater for U251 cells without addition of exogenous GBP-1 (Supp. Figs. 1C).

To further determine how GBP-1 may alter the migration (as measured by wound closure after wounding), the rates of wound closure were determined for each of the 2-hour time increments post wounding (Supp. Fig. 2A). As suggested by the wound closure curves (Supp. Fig. 1B), the rate of closure for U251 cells was faster from zero to 6 hours than the U251+GBP-1 cells. Between 6 and 14 hours the rates of closure were more comparable. The changes of rates of closure over time are visualized graphically first for just the U251 and U251+GBP-1 (D4) cells and then for all three cell lines (Supp. Fig. 2B, C). It is unclear why our U251 cells did not migrate faster in the presence of GBP-1.

### GBP-1 can promote movement of Drp1 to the mitochondria

Cancer cell migration/invasion can be promoted by mitochondrial fission [40,50]. Mitochondrial fission is facilitated by the activation of the normally cytosolic mitochondrial Dynamin-Related Protein 1 (Drp1) and its translocation to the outer membrane of the mitochondria where it drives mitochondrial fission [40,50]. To determine whether GBP-1 promotes movement of Drp1 to mitochondria, subcellular fractionation of U251 cells ± GBP-1 were used to determine the cellular location of Drp1. Drp1 is predominately in the mitochondrial fraction (M) in cells with empty vector but more Drp1 moves from the cytosol (C) to the mitochondria (M) in the presence of elevated GBP-1 (Fig. 3A). In contrast, subcellular fractionation of SNB75 cells showed very little Drp1 associated with the mitochondria (Fig. 3B). The observation of little Drp1 associated with the mitochondria of SNB75 cells is consistent with the long, tubular mitochondria found in these cells (Fig. 1). U251 cells had much smaller mitochondria, suggesting mitochondrial fission was occurring even in the absence of exogenous GBP-1 (Fig. 2).

**Figure 3.**
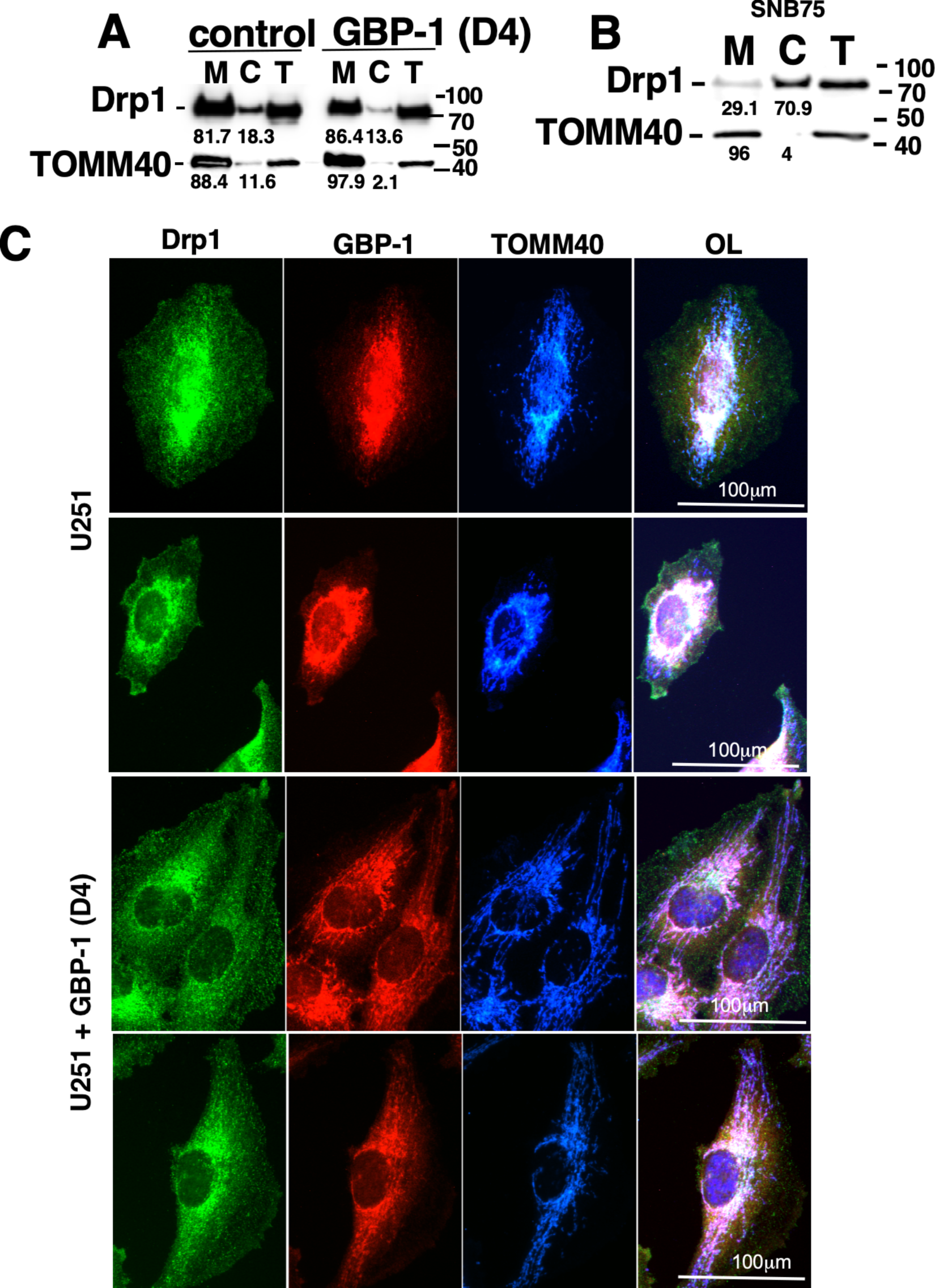
GBP-1 induces the translocation of Drp1 to the mitochondria. **A.** Control (empty vector) and myc GBP-1-expressing U251 cells (D4) were fractionated into mitochondrial (M) and cytosolic fractions (C) and analyzed for Drp1 and TOMM40 by WB. Total cell lysates (T) are also provided. **B.** SNB75 cells were fractionated into mitochondrial (M) and cytosolic fractions (C) and analyzed for Drp1 and TOMM40 by western blot. The numbers under the blots are the percentage of the protein in each fraction. C. Control U251 and U251+GBP-1 (D4) cells were plated on coverslips ON and then analyzed for the expressions of GBP-1, Drp1, and TOMM40 by indirect IF. Two representative examples of each cell type are shown.

To further analyze the intracellular location of Drp1, control U251 or U251+GBP-1 (D4) were analyzed by epifluorescence after IF (Fig. 3C). In the presence or absence of GBP-1, Drp1 distribution was punctate throughout the cytoplasm with some concentration around the nucleus. In the absence of exogenous GBP-1 there is little GBP-1 so the intensity of the images were enhanced to see its localization. While some localization of GBP-1 with mitochondrial TOMM40 is observed, this co-localization is much more visible in the cells with exogenous GBP-1. Some of the Drp1 appears localized to mitochondria in the presence or absence of GBP-1. However, the mitochondrial staining was more spread out throughout the cells in the presence of GBP-1 and close examination suggested association with Drp1 along mitochondrial tracks. This is consistent with the subcellular fractionation (Fig. 3A).

### GBP-1 makes U251 cells less sensitive to inhibition of Drp1

To determine whether migration of U251 cells is sensitive to Drp1, U251 and U251+GBP-1 (D4) cells were analyzed for changes in wound closure in the presence of different concentrations of the Drp1 inhibitor, Mdivi-1 (Supp. Fig. 3). Inhibition of migration of U251 cells (Supp. Fig. 3A) or U251+GBP-1 cells (Supp. Fig. 3B) by Mdivi-1 was dose dependent. The rates of wound closure at the different concentrations of Mdivi-1 were compared (Supp. Fig. 3C). As described previously, the rate of wound closure from 4 to 14 hours post wounding was significantly faster for U251 compared to U251+GBP-1 (D4) cells (Suppl. Fig. 3A and B). While the rate of wound closure for both U251 and U251+GBP-1 cells was inhibited by Mdivi-1, the U251+GBP-1 (D4) showed a smaller decline than the U251 cells (Suppl. Fig. 3C). As stated earlier, the rate of wound closure for untreated U251 cells was significantly shorter than for U251+GBP-1 cells (Suppl. Fig. 1A and 3C). While the wound closure rates of both U251 and U251+GBP-1 cells were shortened by treatment with either 5 or 20 µM of Mdivi-1, the cells without exogenous GBP-1 were more sensitive to Mdivi-1 treatment (Suppl. Fig. 3C). There was no longer a difference in the rate of wound closure between the two cell lines after treatment with either 5 µM or 20 µM drug. The data for 50 µM Mdivi-1 was not included because at this concentration there was evidence of cell toxicity (Supp. Fig. 4) (Note pycnotic nuclei). Consistent with the long tubular mitochondria of SNB 75 cells, little of the Drp1 localized to mitochondria (Fig. 3D) and the cells closed their wound more slowly and exhibited less sensitivity to the Drp1 inhibitor, Mdivi-1 (Supp. Fig. 3D).

### Drp1 co-localizes with GBP-1 at mitochondria

To determine whether Drp1 localizes with GBP-1, U251+GBP-1 (D4) cells were analyzed by triple label IF (Fig. 4). Drp-1 was localized throughout the cytoplasm in a punctate distribution but with concentrations at the inner plasma membrane (denoted by white arrowheads) and within cytosolic regions closer to the nucleus (Fig. 4A). As observed previously, GBP-1 has a punctate distribution within the cytoplasm and at the inner mitochondrial membrane (white arrowheads) (Fig. 4B). However, much of the GBP-1 was associated with mitochondria (Fig. 4B; green arrowhead designates GBP-1 associated with mitochondria in 4C). Close examination showed that the staining of larger structures for GBP-1 line up with the mitochondrial staining for TOMM40. TOMM40 localized closer to the nucleus with smaller mitochondria closer to the plasma membrane (Fig. 4C; green arrowheads show mitochondria co-localizing with GBP-1). Overlays were generated comparing only two of the proteins at a time for easier visualization. Comparing GBP-1 and Drp1 localization again shows that they both have punctate distributions within the cytoplasm and concentrate at the inner plasma membrane (Fig. 4C). Amplification of the region shown in the box in 4C demonstrates that while both Drp1 and GBP-1 went to the inner plasma membrane they exhibit little or no co-localization (as evidenced by yellow color; white arrowhead)(Fig. 4a). Neither did the individual puncta of the proteins in the cytoplasm co-localize (Fig 4a; green arrowhead). Co-localization of the two proteins only occurred at mitochondria (Fig. 4 D,E,a,b,c,d). The localization of the mitochondria varied as fission proceeded. Drp1 localization with mitochondria could be observed at the lamellapodia in some cells (Suppl. Fig. 5).

**Figure 4.**
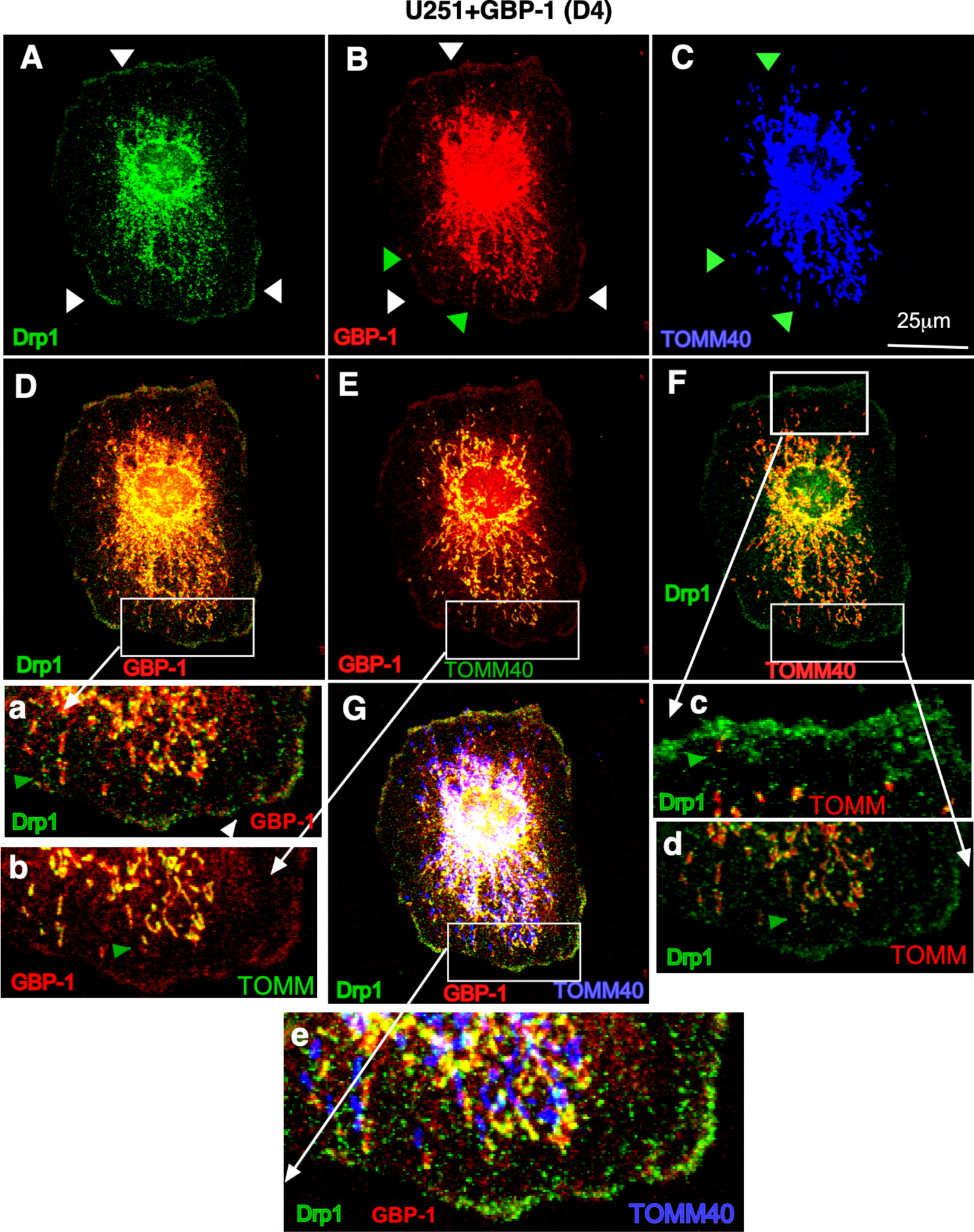
Drp1 colocalizes with GBP-1 at GBM mitochondria. U251 cells expressing myc GBP-1 (D4) were analyzed for the intracellular location of Drp1 (**A**), GBP-1 (**B**), and TOMM40 (**C**) by triple label IF. Images were captured at 0.2 µm z-sections. **D**. The images of Drp1 and GBP-1 staining are overlayed. The white box delineates the region amplified in (**a**). **E**. The images of GBP-1 and TOMM40 are overlayed and the white box delineates the region amplified in (**b**). **F**. The images of Drp1 and TOMM40 are overlayed and the white boxes delineate regions amplified in (**c**) and (**d**). G. The overlay of Drp1, GBP-1 and TOMM40 is shown. The white box delineates the region amplified in (**e**).

### GBP-1 results in shorter, wider mitochondria

Mitochondrial fission is closely correlated with increased cell migration, invasion, lamellipodia formation, and turnover of focal adhesions [40,50,51]. The mitochondria in the U251+GBP-1 cells (D4) were much less filamentous compared to the SNB75 cells (Fig. 1C versus Fig. 2B). To determine if GBP-1 alters mitochondrial dynamics in GBM cells, vector control U251 and U251+GBP-1 cells were stained for TOMM40 and the length and width of individual mitochondrion were measured (Figure 5). Representative figures of the mitochondria are shown (Figure 5A). GBP-1 resulted in mitochondria that were both shorter and slightly wider than those in vector control cells (Figure 5B and C). Their elongation index, calculated from the length divided by the width of individual mitochondria, was also reduced. (Figure 5D). Together these data indicate that GBP-1 results in shorter mitochondria, most likely by promoting fission.

**Figure 5.**
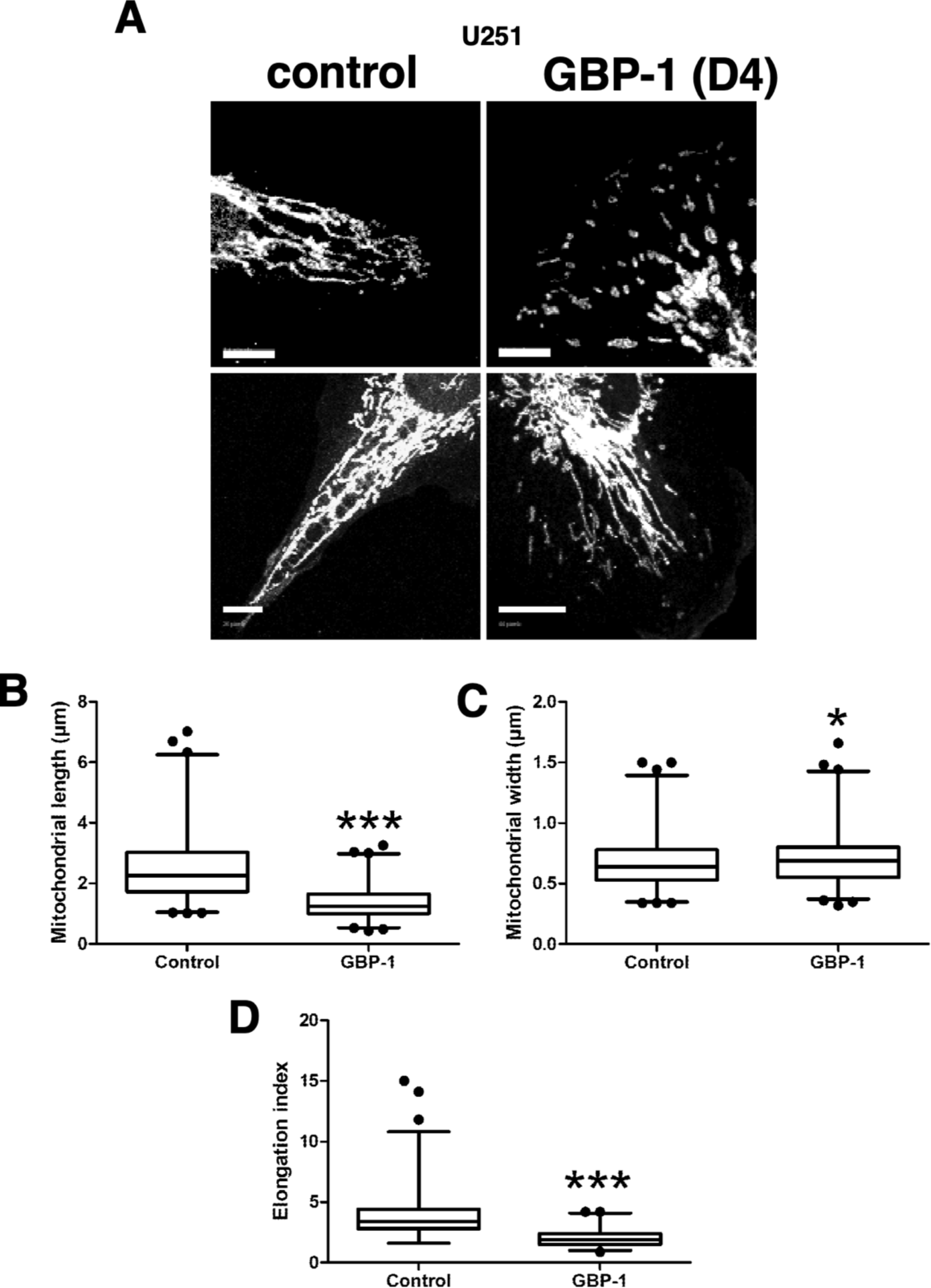
GBP-1 shortens glioblastoma cell mitochondria. U251 cells with (GBP-1) or without (control) myc-epitope tagged GBP-1 were stained for TOMM40 and analyzed by confocal microscopy at 0.2 µm optical sections. **A.** Representative examples of mitochondrial morphology are presented. Size bar = 10 µm. **B.** The length of individual mitochondria were measured as described in Methods. **C.** The width of individual mitochondria was measured as described. **D.** The elongation index of individual mitochondria was calculated by dividing the length by the width. (n = 2; * = p < 0.05; *** = p < 0.001).

## Discussion

The data from several studies of GBMs demonstrate a correlation of GBP-1 with poor prognosis[8,11]. Both overall and progression-free survival of patients with GBM were significantly less with higher expression of GBP-1 [10,11]. GBP-1 expression may be more elevated in GBM with EGFRvIII mutations than with tumors with wild type level or amplification of EGFR [10]. Whether GBP-1 promotes GBM progression by promoting cell migration and/or invasion was analyzed *in vitro* by Boyden chamber using either U251 or SHG44 GBM cells over-expressing GBP-1 [11]. The increase in migration observed was very modest but the increase in invasion was 2-3 fold [11]. Even though the migration increase was modest, multiple genes associated with migration and invasion were increased, such as MMP-9, Twist, and Snail [11]. Intracranial injection of either U251 or SHG44 cells over-expressing GBP-1 resulted in larger tumors and significantly shorter survival times [11]. Conversely, knockdown of GBP-1 in SNB19 cells with intracranial injection resulted in smaller tumor and fewer invasive cells [8].

Our study demonstrates that GBP-1 localizes to GBM mitochondria (Figs. 1, 2, 3). However, our data suggest that GBP-1 at the mitochondria is not always sufficient to promote mitochondrial fission (Figs. 1, 2, 3). GBP-1 localizes to mitochondria in both U251 and SNB75 GBM cells, but only U251 cells are responsive to Drp1 and GBP-1 can further promote movement of Drp1 to the mitochondria (Fig. 3) and act to modulate mitochondria activity. Consistent with this, elevated GBP-1 attenuated the inhibition of U251 cell migration after treatment with the Drp1 inhibitor, Mdivi-1 (Supp. Fig. 3). Fully understanding how GBP-1 enhances mitochondrial fission still requires more investigation. However, there are some properties of GBP-1 that can provide us with potential avenues for investigation. As a member of the dynamin super family, GBP-1 interacts with lipids, forms oligomers, and modifies membranes [18,33,52]. It also modulates both actin [53] and tubulin activity [54,55], both of which are important in mitochondrial fission [56–58]. However, it seems that the earlier steps of mitochondrial fission must occur before GBP-1 can promote further fission.

Both GBPs and Drp1 are GTPases that are members of the Dynamin Superfamily of Proteins (DSPs) [59]. These proteins can be subdivided into a number of subfamilies with different properties and function but there are some common properties of members of the DSPs [52,59–62]. One such common feature is the ability to alter membranes in a GTPase-dependent manner. Family members have a large GTPase domain and differ from many of the GTPases, such as members of the Ras superfamily of GTPases, by exhibiting oligomerization-dependent GTPase activity, low affinity for GTP and GDP, and many exhibit binding to cellular membranes (reviewed in [18,52,59,60,63,64]). Drp1 is a member of a subgroup of DSPs called the dynamin-like proteins (DLPs) that facilitates mitochondrial fission [61,62]. The GBPs are in another subfamily of DSPs with the atlastins. These proteins are not considered DLPs because the only region that is significantly homologous to DLPs is the GTP binding domain of GBPs and even here the region of the G4 homology region is the only DSPs not identical to dynamin. Members of the dynamin family have low affinity for both GTP and GDP. Under normal cellular conditions, the proteins would be expected to be GTP bound. Unlike members of the Ras family of GTPases, members of the DSPs oligomerize, a process that stimulated GTPase activity. This oligomerization dependent activity of the GTPase is usually preceded by membrane binding. The functions of DSPs include vesicle scission, organelle division and fusion, cytokinesis, and antiviral activity.

Mitochondrial dynamics and their disruption play important roles in health and disease; one such disease being cancer [65–67]. Mitochondrial morphology and cellular location are important in determining cellular activities [66]. One such activity that influences mitochondria morphology and cellular location is mitochondrial fission. Mitochondrial fission is important because it generates smaller mitochondria that can be moved by the cytoskeleton to the leading edge of migrating/invading cells where energy is required [68]. Determination of mitochondrial shape and location are governed by complex processes, with many of these that promote mitochondrial fission working through Drp1 [50,69]. In addition to promoting mitochondrial fission, Drp1 may also play a role in lamellipodia formation [70]. As a consequence of enhanced mitochondrial fission, cancer cells exhibit enhanced migration/invasion [40,51,71].

While significant progress has been made in understanding the mechanisms behind mitochondrial fission, there is much that is not yet clear (reviewed in [18,52,59–64,72]). One current model of mitochondrial fission can be divided into 3 steps. The first step involves the actin network of the ER beginning the constriction of mitochondria. This precedes the recruitment of Drp1 to the mitochondria. Drp1, the primary driver of mitochondrial fission, is primarily a cytosolic protein that relocates to the mitochondria when activated [50]. When activated by phosphorylation, Drp1 moves to the mitochondria where it interacts with specific receptors on the outside of the outer membrane [40]. These activated Drp1 molecules are recruited to the pre-constricted regions of the mitochondria where GTP hydrolysis drives further constriction of the mitochondria. Drp1, however, cannot drive the process to the point of fission. The final steps for fission require the recruitment of Dynamin 1 (Dyn2) to the Drp1 already constricted regions. Much remains unclear about the steps involved, the co-factors required, and the cytoplasmic environment required for mitochondrial fission.

The observation that GBP-1 can augment the fission process is intriguing. However, more experimentation is needed to determine how GBP-1 contributes to promoting fission.

Note: While we were examining the consequence of GBP-1 localization to mitochondria in GBM cells, murine GBP-1 (mGBP-1) was shown to regulate dysfunction of mitochondria and senescence in macrophages *in vitro* [73]. mGBP-1 is not confirmed to be the ortholog of human GBP-1. No evidence for a role in mitochondrial fission/fusion and cell migration/invasion was demonstrated in this study. Also, during our study, GBP-2 was demonstrated to inhibit Drp-1 mediated fission in breast cancer cells, which inhibited cancer cell invasion [74,75].

## 5. Conclusions

The expression of GBP-1 and its association with mitochondria in GBM cells promotes the fission of mitochondria, resulting in shorter and wider mitochondria.

## Supplementary Materials

The following supporting information can be downloaded at: www.mdpi.com/xxx/s1, Figure S1: Wound healing for U251 cells with or without exogenous GBP-1; Figure S2: Rates of wound closure for U251 cells with or without exogenous GBP-1; Figure S3: Mdivi-1 sensitivity of U251 cells with or without exogenous GBP-1; Figure S4: Representative images of 50 µM Mdivi-1 treated cells; Figure S5: Localization of Drp1 and TOMM40 in U251 cells plus GBP-1.

## Author Contributions

Conceptualization D.J.V., A.L.N-K.; methodology D.J.V., A.L.N-K.;, M.P.M., G.O.N.; validation S.M., M.P.M., A.L.N-K., D.J.V.; formal analysis D.J.V., M.P.M., A.L.N-K.; investigation R.J.K., G.O.N., M.P.M., S.M., J.S.J., A.L.N-K., D.J.V.; writing—original draft preparation D.J.V.; writing—review and editing, M.P.M., D.J.V., G.O.N., R.J.K., S.M.; A.L.N-K., J.S.J.; visualization D.J.V., A.L.N-K., M.P.M., R.J.K., J.S.J.; supervision D.J.V., A.L.N-K., project administration D.J.V., funding acquisition D.J.V., A.L.N-K.. All authors have read and agreed to the published version of the manuscript.

## Data Availability Statement

Most of the data generated from this study are included in this article (and the Supplementary Data). The files containing the images used to measure the mitochondria can be made available upon request.

## Acknowledgments

The authors thank Stephanie Angel for the generation of U251 cells containing myc-hGBP-1. The authors also thank Dr. William Maltese, Dr. Katherine Eisenmann, Dr. Jean Overmeyer, and Kristi Pettee for helpful discussions about glioblastoma and their culturing.

## Footnotes

1 Our results differ from those in reference [8] where GBP1 was expressed prior to any treatment in U87, U178, U373, LN308, T98G, LN229, LN235, U251, A1207, and DBTRG cells. Please note that GBP-1 is also cytokine-induced and part of innate and adaptive immunity. A negative control for the previous study would have been helpful.

