## Supplemental Figures for "The large GTPase Guanylate-Binding Protein-1 (GBP-1) promotes mitochondrial fission in glioblastoma"

**Figure Legends for Supplementary Figures.**

**Supplementary Figure 1. A.** Wound healing was performed as described on U251 cells, U251 cells + GBP-1 (D4), and U251+ GBP-1 (D7) cell lines. Each line on the graph represents the data from measurements of 5 wells from one experiment. The numbers 1 and 2 for each cell line represent the two replicates. **B.** Representative images of wound closures for U251 and U251+GBP-1 (D4) and U251+GBP-1 (D7) cells for 4 and 14 hours after wounding are shown. **C.** The average rate of wound closure was shown for all three cell lines. The averages were determined by calculating the slope of the line including the times from 4 hours through 14 hours. The data was obtained from 2 experiments measuring 5 wells per experiment. Data is presented as average slope ± SEM. *** = $p < 0.001$, ** = $p < 0.005$.

**Supplementary Figure 2. A.** The rates of wound closure for each of the 2-hour intervals post wounding were determined from the slope of the line between the time points. The data in red illustrates that the rate of closure for these time points were faster than observed for the other cells. **B.** The changes in rate of wound healing for the 2-hour time intervals were graphed for each experiment of U251 and U251+ GBP-1 (D4) cells. **C.** The changes in rate of wound healing for the 2 hour time intervals were graphed for each experiment of U251, U251+GBP-1 (D4), U251+ GBP-1 (D7).

**Supplementary Figure 3. A.** U251 cells were grown to confluence, wounded, and analyzed for wound closure over time in the presence of the concentrations of Drp1 inhibitor, Mdivi-1, listed. The data of a representative experiment is shown. **B**. U251 cells expressing GBP-1 (D4) were grown to confluence, wounded, and analyzed for wound closure over time in the presence of the concentrations of Drp1 inhibitor, Mdivi-1, listed. The data of a representative experiment is shown. **C.** The average rates of wound closure of U251 cells and U251 cells expressing GBP-1 treated with different concentrations of Mdivi-1 were determined as described in Methods for the time points from 4 hours to 14 hours post wounding. Data was presented for 5 wells each for 2 experiments as average ± SEM. * = $p < 0.05$, ** = $p < 0.005$, *** = $p < 0.001$, n.s. = not significant. **D.** SNB75 cells were grown to confluence, wounded, and analyzed for wound closure over time in the presence of the concentrations of Drp1 inhibitor, Mdivi-1, listed. The data of a representative experiment is shown.

**Supplementary Figure 4.** Representative images of wound closures for U251 and U251+GBP-1 (D4) treated with 50 μM Mdivi-1 for the times listed post wounding.

**Supplementary Figure 5.** U251+GBP-1 cells were stained for Drp1, TOMM40 and DAPI and analyzed by multiphoton confocal microscopy at 0.5 μm optical sections. Z-stacks were compiled and representative examples of the distribution of Drp1 and TOMM40 are shown. The area denoted with the red rectangle is enlarged and the individual channels from that region are shown on the left. Arrowheads point toward other areas of the cells were TOMM40 and Drp1 co-localize at the plasma membrane.

### Supplementary Figure 1

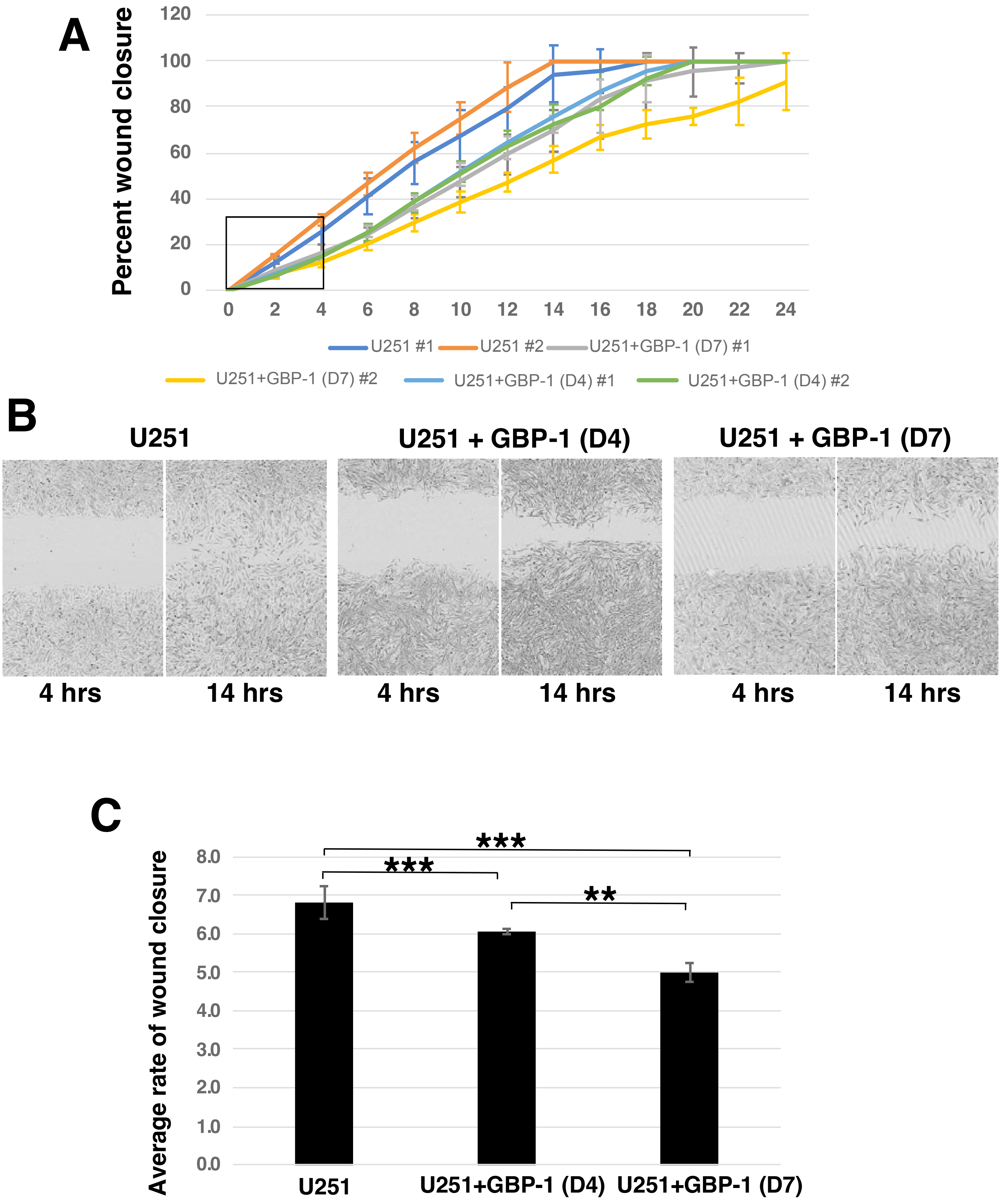

### Supplementary Figure 2

**A**

#### Rate of Wound Closure

Hours post wounding

| | U251 #1 | U251 #2 | D4 #1 | D4 #2 | D7 #1 | D7 #2 |
|---|---|---|---|---|---|---|
| 0 to 2 | 5.85 | 7.4 | 3.64 | 3.33 | 4.52 | 3.44 |
| 2 to 4 | 6.92 | 7.92 | 3.73 | 4.03 | 3.45 | 2.74 |
| 4 to 6 | 7.61 | 7.87 | 5.34 | 5.5 | 4.08 | 3.99 |
| 6 to 8 | 7.4 | 7.77 | 6.42 | 6.26 | 5.84 | 4.65 |
| 8 to 10 | 6 | 6.36 | 6.77 | 6.32 | 5.84 | 4.51 |
| 10 to 12 | 5.86 | 6.83 | 6.36 | 5.69 | 5.86 | 4.26 |
| 12 to 14 | 7.53 | 5.76 | 5.62 | 5.19 | 5.22 | 4.93 |
| 14 to 16 | 0.68 | 0 | 5.52 | 3.83 | 6.83 | 4.73 |
| 16 to 18 | 2.15 | 0 | 4.34 | 6.09 | 4.07 | 2.86 |
| 18 h to 20 h | 0 | 0 | 2.27 | 3.77 | 1.03 | 1.95 |
| 20 h to 22 h | 0 | 0 | 0 | 0 | 0.8 | 3.05 |
| 22 h to 24 h | 0 | 0 | 0 | 0 | 1.58 | 4.43 |

**B**

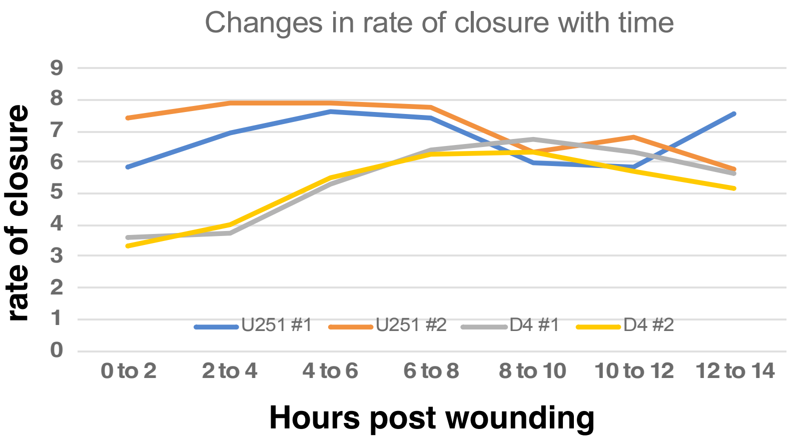


**C**

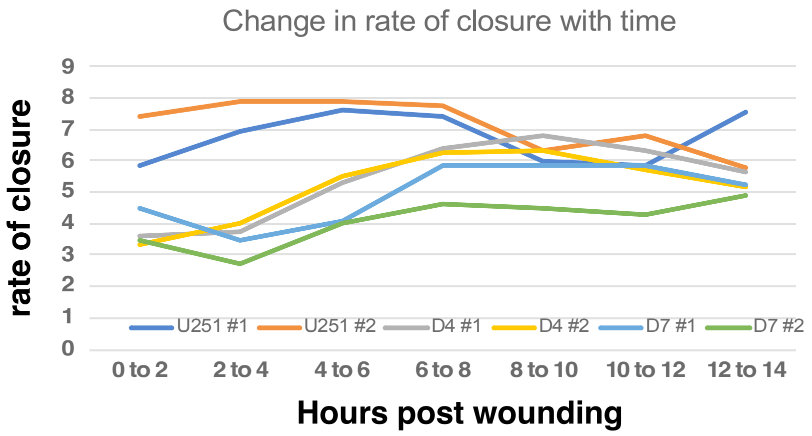

### Supplementary Figure 3

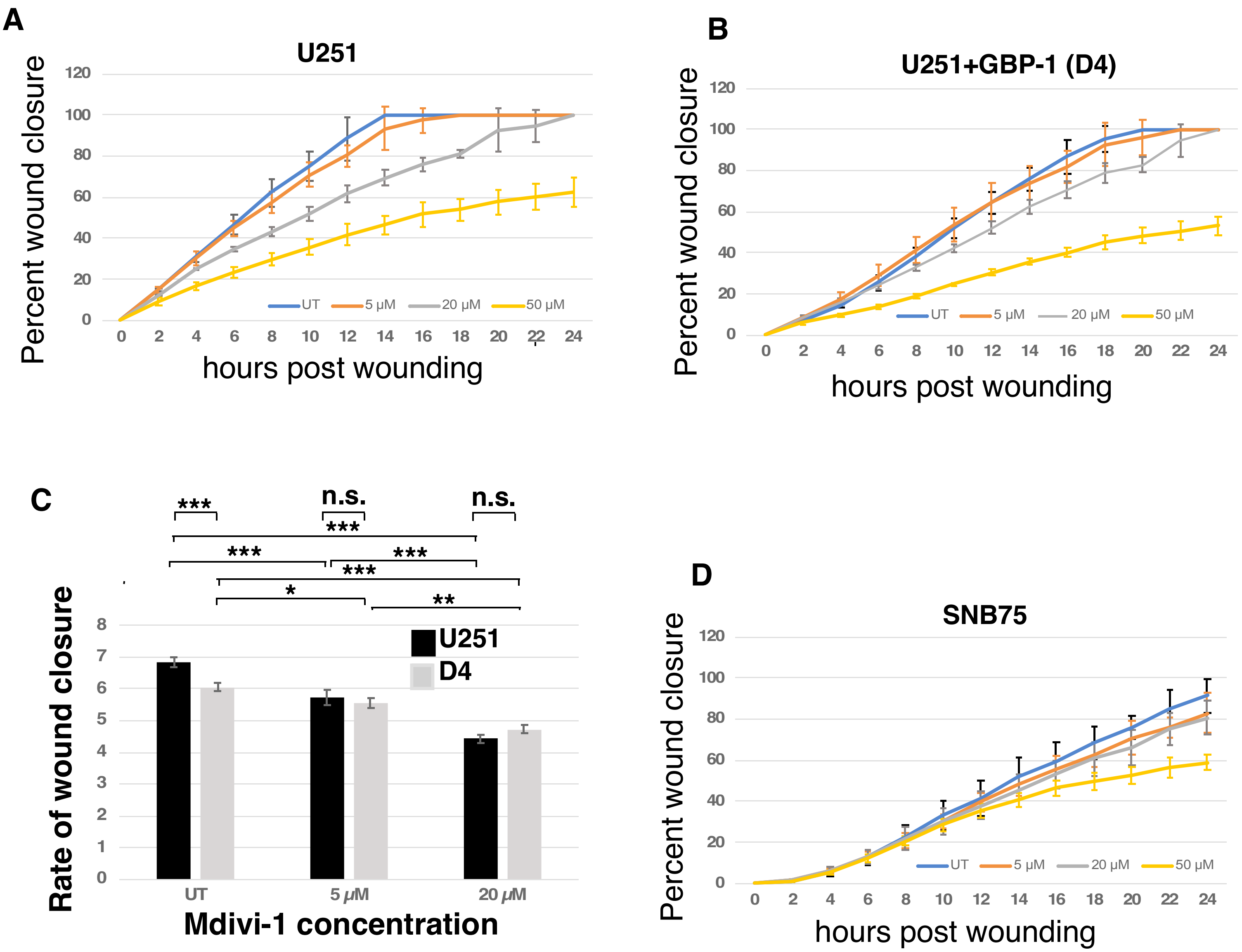

### Supplementary Figure 4

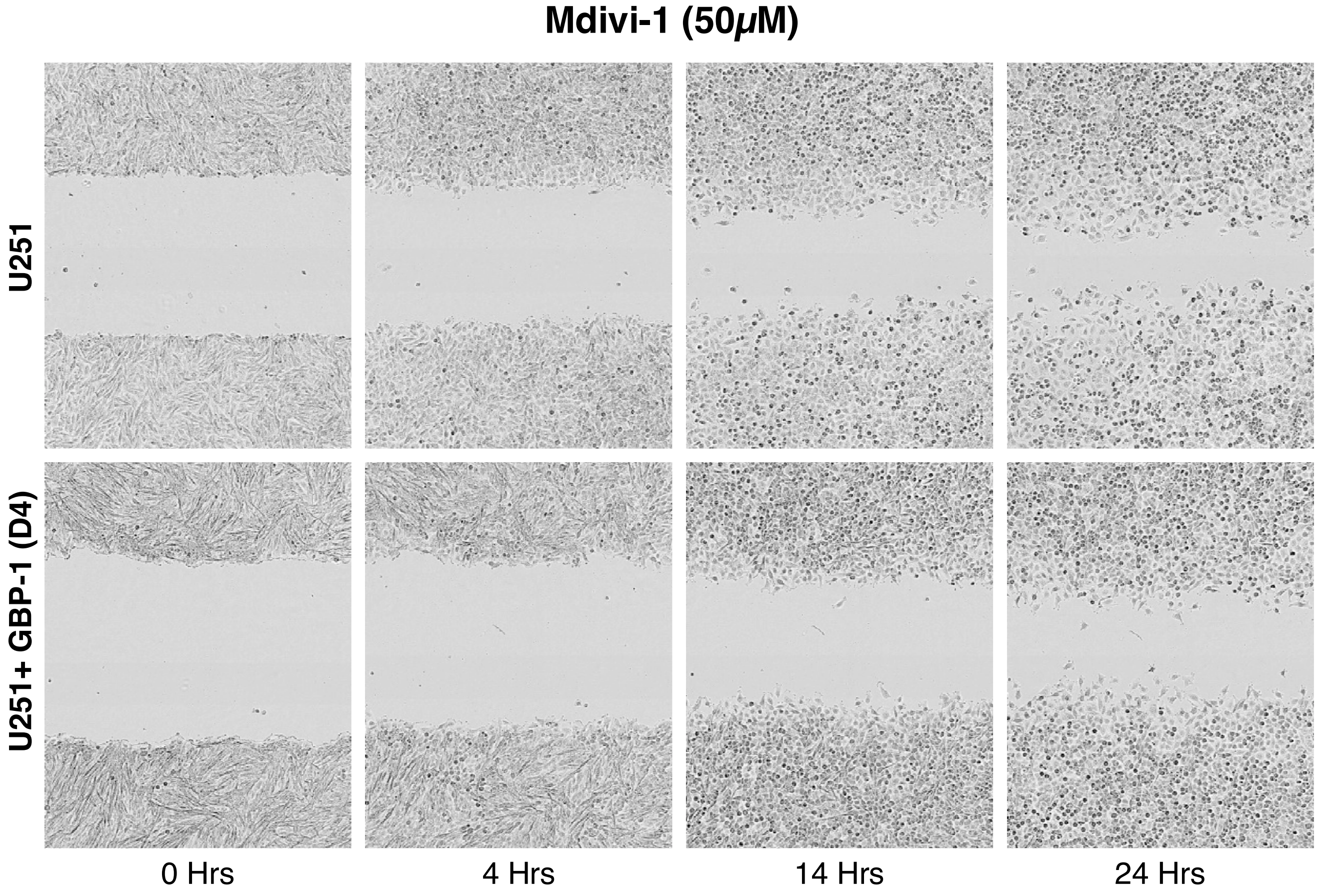

### Supplementary Figure 5

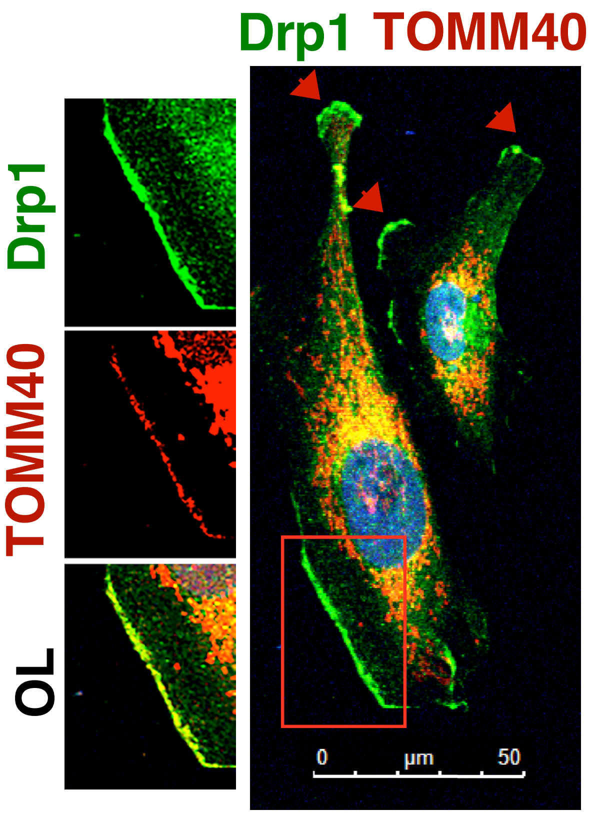
